# Upper Airway Gene Expression in Hospitalized Children with Rhinovirus-induced Respiratory Illnesses

**DOI:** 10.1101/2025.04.29.651288

**Authors:** Jordan E. Kreger, Alex L. Sliwicki, Saly N. Essoh, Yiran Li, Chaandini Jayachandran, Jessica A. Czapla, Toby C. Lewis, Erin M. Kirkham, Rodney J. Vergotine, Antonia P. Popova, Heidi R. Flori, Marc B. Hershenson

**Affiliations:** Departments of Pediatrics, University of Michigan Medical School, Ann Arbor, MI; Departments of Otolaryngology, University of Michigan Medical School, Ann Arbor, MI; Departments of Molecular and Integrative Physiology, University of Michigan Medical School, Ann Arbor, MI; Department of Orthodontics and Pediatric Dentistry, School of Dentistry, Ann Arbor, MI

**Author notes:** Corresponding author: Marc B. Hershenson, Medical Sciences Research Building II, 1150 W. Medical Center Drive, Ann Arbor, MI; e mail. Funding: This work was supported NIH grants K23HL153897 (E.M.K.), R01HL167716 (A.P.P.), R01HL149910 (H.R.F.), R01AI120526, R56AI150660 and R01AI155444 (M.B.H.). **Conflict of Interest Statement:** The authors certify that they have NO affiliations with or involvement in any organization or entity with any financial interest (such as honoraria; educational grants; participation in speakers’ bureaus; membership, employment, consultancies, stock ownership, or other equity interest; and expert testimony or patent-licensing arrangements), or non-financial interest (such as personal or professional relationships, affiliations, knowledge or beliefs) in the subject matter or materials discussed in this manuscript.

**Keywords:** asthma, bronchiolitis, children, chemotaxis, cilia, cysteinyl leukotrienes, epithelial-mesenchymal transition, goblet cells, mast cell, respiratory syncytial virus, rhinovirus

## Abstract

**Background:** The precise mechanisms underlying rhinovirus (RV)-induced respiratory illnesses are not completely known.

**Objective:** We sought to obtain nasal transcriptomic data from hospitalized children with respiratory viral infections.

**Methods:** We obtained nasal swabs from 46 children with RV (16 RV-A, 30 RV-C). For comparison, we examined swabs from 12 children with RSV and six controls. Subjects ranged in age from 1 month to 18 years. Viral detection, genotyping and copy number were determined by PCR. RNA transcripts were measured by next generation sequencing and differences in gene expression calculated using DESeq2.

**Results:** Compared to controls, 1232 transcripts were upregulated (adjusted p<0.05, fold change >1.5) by all three viruses, including genes regulating granulocyte chemotaxis, cysteinyl leukotriene production, epithelial remodeling and antiviral responses. Cilium-related genes were downregulated. Compared to RSV, RV induced greater expression of 207 genes including those regulating eosinophilic inflammation, mucus secretion and mast cell function.RSV induced greater upregulation of 674 genes including those regulating neutrophilic inflammation and type 1 IFN response. Computational deconvolution of RNA-seq profiles revealed that viral infection decreased ciliated cells while increasing neutrophils, natural killer cells, monocytes (all viral species) and goblet cells (RV only). RV-C infections increased mast cells and IFN-λ mRNA expression. RV copy number correlated with the expression of mast cell proteases and numerous pro-inflammatory and IFN-stimulated genes.

**Conclusion:** Children hospitalized with RV and RSV infections mount robust inflammatory responses, but virus-specific differences exist.

**Clinical Implication:** These data provide insight into mechanisms by which RV, and in particular, RV-C, trigger respiratory illnesses.

**Capsule summary:** Nasal transcriptomics demonstrate that RV infections in hospitalized children induce expression of genes regulating eosinophilic inflammation, mucus secretion and mast cell function, with RV-C in particular increasing IFN-λ expression.

## Introduction

Rhinovirus (RV) is a small picornavirus grouped into the genus *Enterovirus*. RV has an icosahedral, non-enveloped viral capsid carrying a positive sense, single-stranded RNA genome of ∼7,200 bp ^1, 2^. Analysis of the over 100 RV serotypes collected in the 1960s divided them into two species on the basis of susceptibility to antivirals: RV-A and RV-B ^3^. RV strains were also classified by receptor specificity, either intercellular adhesion molecule-1 (ICAM-1) (the major group) or low-density lipoprotein receptor (LDLR) (the minor group) ^4, 5^. Subsequently, untypeable RVs that failed to produce cytopathic effects in HeLa cell culture were described using PCR. First reported in 2007 ^6, 7^, RV-C has been associated with severe respiratory illnesses in children and adults, including wheezing, O_2_ supplementation, hospitalization and ICU admission ^8–18^. RV-C has been linked to bronchiolitis in young children ^19, 20^. Infections with RV-C are more likely to occur in children with a history of asthma or who develop asthma ^11, 14–17^. Not all studies have found differences in clinical outcome between different RV species, however ^19, 21, 22^. Thus far, 55 RV-C genotypes have been reported ^23, 24^. In contrast to RV-A and RV-B, RV-C utilizes cadherin-related family member 3 (CDHR3) as a receptor ^25^. CDHR3 is most highly expressed in immature ciliated airway epithelial cells ^26^ and localized to acetylated α-tubulin-positive cilia ^27^.

The mechanisms underlying RV-induced respiratory illnesses have been an area of active study for many years. RV infection induces the release of chemokines from the airway epithelium, attracting pro-inflammatory neutrophils ^28–30^, lymphocytes ^31^, eosinophils ^32, 33^ and type 2 innate lymphoid cells ^34^. However, nearly all human studies have employed patients undergoing experimental RV infection. Leveraging the concordance in gene expression between upper and lower airway samples ^35^, researchers have examined the nasal transcriptome of children with respiratory viral infections ^36–41^. A multicenter, prospective, longitudinal study of asthma exacerbations in 6-17 year-old children ^40^ identified gene expression modules including upregulation of SMAD3 signaling, extracellular matrix production, mucus secretion, eosinophil activation and type I interferon (IFN) response.

Numerous questions about viral-induced respiratory illnesses remain. For example, we have shown in a cell culture model of injured/regenerating airway epithelium that RV infection induces epithelial-to-mesenchymal transition (EMT)-like features such as reduced transepithelial resistance and reduced expression of occludin and E-cadherin ^42^. However, evidence for epithelial remodeling in patients with respiratory viral infections is lacking. Also, while mast cells ^43^ and expression of mast cell proteases ^44^ have been shown to be increased in adults with asthma, the contribution of mast cells to viral-induced respiratory illnesses in children has not been studied.

In this study, we collected nasal swabs from infants and children hospitalized with RV infection and assessed gene expression by next generation sequencing, comparing profiles to swabs obtained from control subjects undergoing otolaryngologic procedures. We also compared gene expression patterns to those obtained from children with respiratory syncytial virus (RSV) infection.

## Methods

### Patient selection and data collection

This study was approved by the University of Michigan IRB (ID# HUM00133414). Patient recruitment occurred from March 2023 to March 2024. Children hospitalized on a pediatric ward or pediatric intensive care unit with a positive respiratory panel (Biofire FilmArray, Salt Lake City, UT) for enterovirus or RSV infection were identified from the electronic medical record and approached for informed consent. We excluded patients being treated with antibiotics who were judged by the intensive care unit staff to have bacterial pneumonia. For comparison, we also obtained nasal swabs from children undergoing otolaryngologic procedures in the operating room. Virus-infected patients ranged in age from 1 month to 18 years; 26 children were less than two years of age, and 32 were greater than 2 years of age (Table I).

**Table I.** Patient demographics.

|  | Total<br>virus-positive<br>(n=58) | RV-A<br>(n=16) | RV-C<br>(n=30) | RSV<br>(n=12) | control<br>(n=6) | p value* |
| --- | --- | --- | --- | --- | --- | --- |
| Age at consent, mos |  |  |  |  |  | 0.0014 |
| Median | 33.0 | 63.5 | 44.5 | 5.5 | 148.0 |  |
| IQR | 61.5 | 70.5 | 47.8 | 25.3 | 86.8 |  |
| Range | 2-219 | 6-170 | 10-146 | 2-219 | 44-200 |  |
| Gestational age, wks |  |  |  |  |  | 0.3597 |
| Median | 39.1 | 39.6 | 39.1 | 37.9 | 40.0 |  |
| IQR | 3.61 | 1.14 | 6.00 | 2.53 | 0.71 |  |
| Range | 23.7-41.1 | 30.6-41.1 | 24.3-41.0 | 23.7-40.7 | 35.7-40.0 |  |
| Sex, n (%) |  |  |  |  |  | 0.4103 |
| Male | 34 (58.6) | 10 (62.5) | 19 (63.3) | 5 (41.7) | 5 (83.3) |  |
| Female | 24 (41.4) | 6 (37.5) | 11 (36.7) | 7 (58.3) | 1 (16.7) |  |
| Race, n (%) |  |  |  |  |  | 0.6974 |
| White | 33 (56.9) | 8 (50.0) | 17 (56.7) | 8 (66.7) | 4 (66.7) |  |
| Black | 12 (20.7) | 3 (18.8) | 7 (23.3) | 2 (16.7) | 0 (0.0) |  |
| Asian | 3 (5.2) | 0 (0.0) | 2 (6.7) | 1 (8.3) | 0 (0.0) |  |
| Middle Eastern | 1 (1.7) | 0 (0.0) | 1 (3.3) | 0 (0.0) | 0 (0.0) |  |
| Pacific Islander | 1 (1.7) | 1 (6.3) | 0 (0.0) | 0 (0.0) | 0 (0.0) |  |
| More than One Race | 7 (12.1) | 4 (25.0) | 2 (6.7) | 1 (8.3) | 1 (16.7) |  |
| Unspecified | 1 (1.7) | 0 (0.0) | 1 (3.3) | 0 (0.0) | 1 (16.7) |  |
| Ethnicity, n (%) |  |  |  |  |  | 0.9220 |
| Hispanic | 6 (10.3) | 1 (6.3) | 4 (13.3) | 1 (8.3) | 0 (0.0) |  |
| Not Hispanic | 52 (89.7) | 15 (93.8) | 26 (86.7) | 11 (91.7) | 6 (100) |  |
| Level of care, n (%) |  |  |  |  |  | 0.2730 |
| Pediatric ICU | 20 (34.5) | 8 (50.0) | 8 (26.7) | 4 (33.3) |  |  |
| General inpatient | 38 (65.5) | 8 (50.0) | 22 (73.3) | 8 (66.7) |  |  |
| Day of Admission |  |  |  |  |  | 0.3808 |
| Median | 3 | 2 | 2 | 3 |  |  |
| IQR | 1.0 | 1.0 | 1.0 | 0.3 |  |  |
| Range | 1-5 | 1-5 | 1-4 | 2-5 |  |  |
| Admission diagnosis, n (%)† |  |  |  |  |  | 0.0012 |
| Status asthmaticus | 28 (48.3) | 9 (56.3) | 18 (60.0) | 1 (8.3) |  |  |
| Bronchiolitis | 10 (17.2) | 1 (6.3) | 2 (6.7) | 7 (58.3) |  |  |
| Acute respiratory failure | 20 (34.5) | 6 (37.5) | 10 (33.3) | 4 (33.3) |  |  |
| Persistent asthma, n (%)‡ | 19 (32.8) | 7 (43.8) | 11 (36.7) | 1 (8.3) |  | 0.1267 |

\* Statistical significance was assessed by the Kruskal-Wallis test for non-parametric data or Fisher's exact test, where applicable.
† Status asthmaticus includes 'reactive airways disease with exacerbation.' Acute respiratory failure includes 'increased work of breathing,' 'respiratory distress' and 'hypoxia.'
‡ Persistent asthma indicates a physician diagnosis of asthma on controller medication (includes two RV-C patients with bronchopulmonary dysplasia).

**Table II.**
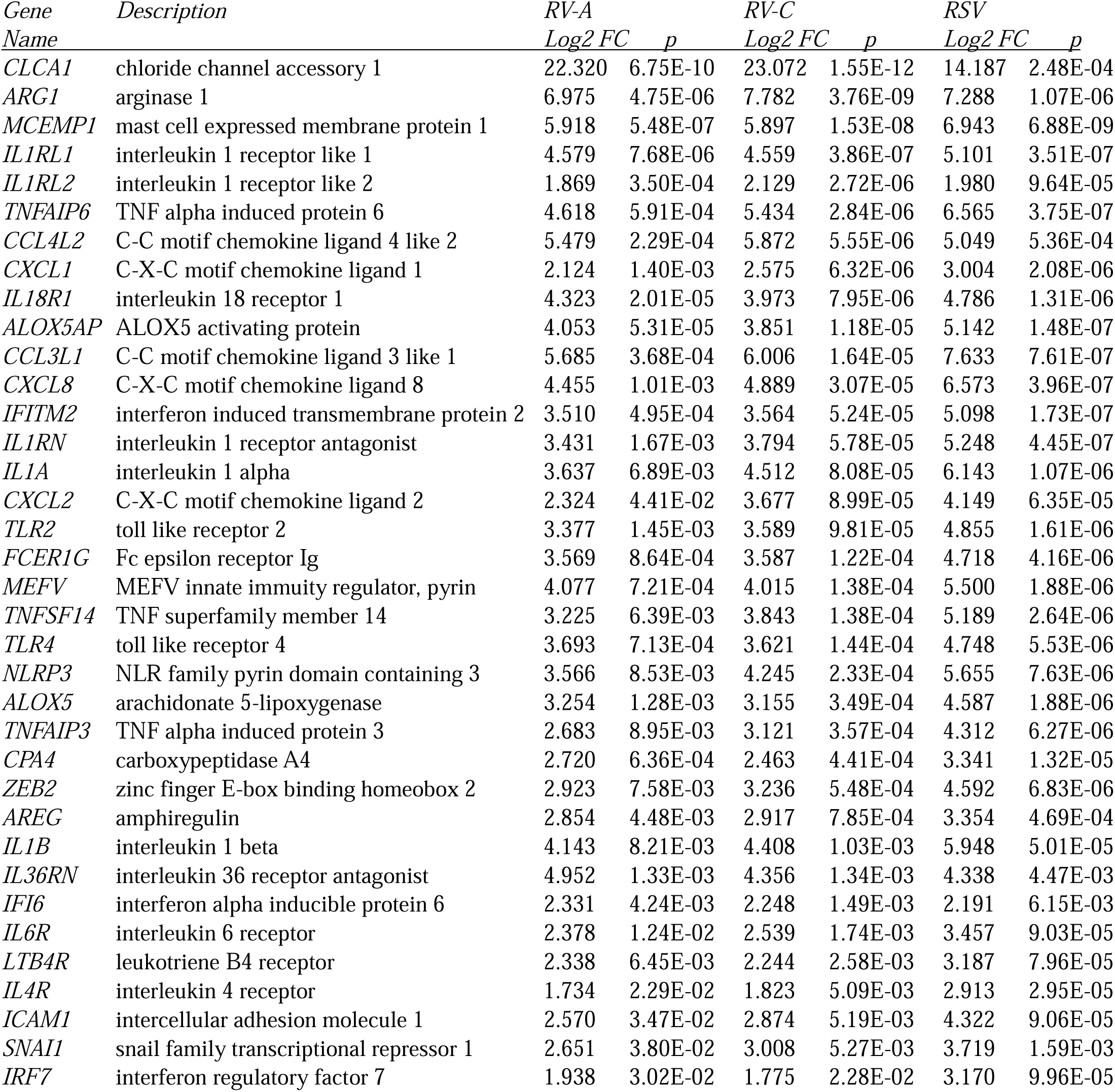
Selected genes significantly upregulated by all three viral species, as determined using DESeq2.

| *Gene*<br>*Name* | *Description* | *RV-A* |  | *RV-C* |  | *RSV* |  |
| --- | --- | --- | --- | --- | --- | --- | --- |
|  |  | *Log2 FC* | *p* | *Log2 FC* | *p* | *Log2 FC* | *p* |
| *CLCA1* | chloride channel accessory 1 | 22.320 | 6.75E-10 | 23.072 | 1.55E-12 | 14.187 | 2.48E-04 |
| *ARG1* | arginase 1 | 6.975 | 4.75E-06 | 7.782 | 3.76E-09 | 7.288 | 1.07E-06 |
| *MCEMP1* | mast cell expressed membrane protein 1 | 5.918 | 5.48E-07 | 5.897 | 1.53E-08 | 6.943 | 6.88E-09 |
| *IL1RL1* | interleukin 1 receptor like 1 | 4.579 | 7.68E-06 | 4.559 | 3.86E-07 | 5.101 | 3.51E-07 |
| *IL1RL2* | interleukin 1 receptor like 2 | 1.869 | 3.50E-04 | 2.129 | 2.72E-06 | 1.980 | 9.64E-05 |
| *TNFAIP6* | TNF alpha induced protein 6 | 4.618 | 5.91E-04 | 5.434 | 2.84E-06 | 6.565 | 3.75E-07 |
| *CCL4L2* | C-C motif chemokine ligand 4 like 2 | 5.479 | 2.29E-04 | 5.872 | 5.55E-06 | 5.049 | 5.36E-04 |
| *CXCL1* | C-X-C motif chemokine ligand 1 | 2.124 | 1.40E-03 | 2.575 | 6.32E-06 | 3.004 | 2.08E-06 |
| *IL18R1* | interleukin 18 receptor 1 | 4.323 | 2.01E-05 | 3.973 | 7.95E-06 | 4.786 | 1.31E-06 |
| *ALOX5AP* | ALOX5 activating protein | 4.053 | 5.31E-05 | 3.851 | 1.18E-05 | 5.142 | 1.48E-07 |
| *CCL3L1* | C-C motif chemokine ligand 3 like 1 | 5.685 | 3.68E-04 | 6.006 | 1.64E-05 | 7.633 | 7.61E-07 |
| *CXCL8* | C-X-C motif chemokine ligand 8 | 4.455 | 1.01E-03 | 4.889 | 3.07E-05 | 6.573 | 3.96E-07 |
| *IFITM2* | interferon induced transmembrane protein 2 | 3.510 | 4.95E-04 | 3.564 | 5.24E-05 | 5.098 | 1.73E-07 |
| *IL1RN* | interleukin 1 receptor antagonist | 3.431 | 1.67E-03 | 3.794 | 5.78E-05 | 5.248 | 4.45E-07 |
| *IL1A* | interleukin 1 alpha | 3.637 | 6.89E-03 | 4.512 | 8.08E-05 | 6.143 | 1.07E-06 |
| *CXCL2* | C-X-C motif chemokine ligand 2 | 2.324 | 4.41E-02 | 3.677 | 8.99E-05 | 4.149 | 6.35E-05 |
| *TLR2* | toll like receptor 2 | 3.377 | 1.45E-03 | 3.589 | 9.81E-05 | 4.855 | 1.61E-06 |
| *FCER1G* | Fc epsilon receptor Ig | 3.569 | 8.64E-04 | 3.587 | 1.22E-04 | 4.718 | 4.16E-06 |
| *MEFV* | MEFV innate immunity regulator, pyrin | 4.077 | 7.21E-04 | 4.015 | 1.38E-04 | 5.500 | 1.88E-06 |
| *TNFSF14* | TNF superfamily member 14 | 3.225 | 6.39E-03 | 3.843 | 1.38E-04 | 5.189 | 2.64E-06 |
| *TLR4* | toll like receptor 4 | 3.693 | 7.13E-04 | 3.621 | 1.44E-04 | 4.748 | 5.53E-06 |
| *NLRP3* | NLR family pyrin domain containing 3 | 3.566 | 8.53E-03 | 4.245 | 2.33E-04 | 5.655 | 7.63E-06 |
| *ALOX5* | arachidonate 5-lipoxygenase | 3.254 | 1.28E-03 | 3.155 | 3.49E-04 | 4.587 | 1.88E-06 |
| *TNFAIP3* | TNF alpha induced protein 3 | 2.683 | 8.95E-03 | 3.121 | 3.57E-04 | 4.312 | 6.27E-06 |
| *CPA4* | carboxypeptidase A4 | 2.720 | 6.36E-04 | 2.463 | 4.41E-04 | 3.341 | 1.32E-05 |
| *ZEB2* | zinc finger E-box binding homeobox 2 | 2.923 | 7.58E-03 | 3.236 | 5.48E-04 | 4.592 | 6.83E-06 |
| *AREG* | amphiregulin | 2.854 | 4.48E-03 | 2.917 | 7.85E-04 | 3.354 | 4.69E-04 |
| *IL1B* | interleukin 1 beta | 4.143 | 8.21E-03 | 4.408 | 1.03E-03 | 5.948 | 5.01E-05 |
| *IL36RN* | interleukin 36 receptor antagonist | 4.952 | 1.33E-03 | 4.356 | 1.34E-03 | 4.338 | 4.47E-03 |
| *IFI6* | interferon alpha inducible protein 6 | 2.331 | 4.24E-03 | 2.248 | 1.49E-03 | 2.191 | 6.15E-03 |
| *IL6R* | interleukin 6 receptor | 2.378 | 1.24E-02 | 2.539 | 1.74E-03 | 3.457 | 9.03E-05 |
| *LTB4R* | leukotriene B4 receptor | 2.338 | 6.45E-03 | 2.244 | 2.58E-03 | 3.187 | 7.96E-05 |
| *IL4R* | interleukin 4 receptor | 1.734 | 2.29E-02 | 1.823 | 5.09E-03 | 2.913 | 2.95E-05 |
| *ICAM1* | intercellular adhesion molecule 1 | 2.570 | 3.47E-02 | 2.874 | 5.19E-03 | 4.322 | 9.06E-05 |
| *SNAI1* | snail family transcriptional repressor 1 | 2.651 | 3.80E-02 | 3.008 | 5.27E-03 | 3.719 | 1.59E-03 |
| *IRF7* | interferon regulatory factor 7 | 1.938 | 3.02E-02 | 1.775 | 2.28E-02 | 3.170 | 9.96E-05 |

### Nasal swabs

Nasal swab samples from hospitalized patients were collected as soon as it was considered safe to do so by the medical team. Samples were taken from 1-5 days after admission (median, 3 days). Samples were taken from the surface of the nasal inferior turbinate using a Copan floxed swab (Murrieta, CA), placed in RNAlater (ThermoFisher Scientific, Waltham, MA), and brought to the laboratory for processing and analysis.

### RV typing

Total RNA was isolated through TRIzol extraction (ThermoFisher) and purified using an RNEasy Mini kit with DNase incubation (Qiagen, Valencia, CA). First-strand cDNA was prepared using the High-Capacity cDNA Reverse Transcriptase kit (ThermoFisher) according to the manufacturer protocol. The P1-P2 region was amplified using semi-nested polymerase chain reaction (PCR) ^8^, purified on an agarose gel, and extracted using a QIAquick kit (Qiagen) for RV typing. Sanger sequencing was performed by Genewiz (South Plainfield, NJ). The identity of each sequence was determined by comparison to known 5′ sequences using NCBI BLAST (http://blast.ncbi.nlm.nih.gov/Blast.cgi).

### Viral detection

For control subjects and patients with RSV-positive swabs, we ruled out asymptomatic viral infection or incidental viral co-infections by analyzing samples using the Seegene Novaplex Respiratory Panels 1A, 2, and 3 (Irvine, CA). The Seegene kit detects human adenovirus, bocavirus 1-4, coronaviruses 229E/NL63 and OC43, enterovirus, influenza A and B, metapneumovirus, parainfluenza viruses 1-4, RSV A and B, and RV-A, -B and -C by multiplex qPCR.

### RV quantification

For samples testing positive for RV, copy number was determined by qPCR. We designed pan-RV primers based on the GenBank sequences of RV-C15 W10 (accession number GU219984.1), RV-A1B (GCA_008798635.1) and RV-A16 (LC699418.1) (Table E1). Plasmids encoding full-length RV-A16 ^45^ and RV-C15 ^46^ were used as copy number standards. Plasmids were provided by Drs. W-M. Lee and Y. Bochkov (University of Wisconsin), respectively.

### Quantitative PCR

Changes in the expression level of selected genes encoding type 2 cytokines and IFNs with very low numbers of transcripts were assessed by qPCR. Oligonucleotide primers used for qPCR are shown in Table E1 in the Online Repository.

Samples were diluted to the same input RNA concentration, cDNA was generated as described above, and results were normalized to 1 µg of total RNA input.

### Statistical analysis

When comparing group differences, statistical significance was assessed by the Kruskal-Wallis test with correction for multiple comparisons by the two-stage linear step-up procedure of Benjamini, Krieger, and Yekutieli for non-parametric, continuous variables, or the Fisher’s exact test for discrete variables. The significance of correlations between viral copy and gene expression was determined using the Pearson correlation coefficient. A Pearson correlation of 0.5 was considered a positive correlation and a coefficient value between ± 0.40 and ± 0.49 was considered a moderate correlation.

### Next generation RNA sequencing (RNA-Seq)

Total RNA was prepared as described above. Measurement of RNA concentration and RIN (RNA integrity number), library generation and next generation sequencing were performed by the University of Michigan Advanced Genomic Core. RIN was determined using the Agilent RNA ScreenTape assay (Santa Clara, CA). Samples of sufficient RNA quality were analyzed by RNA-Seq. Of note, sequencing was conducted on three separate occasions; to control for batch effects, technical replicates were included in each group. cDNA libraries for next generation sequencing were prepared using the SMART-Seq v4 PLUS kit (Takara, San Jose, CA). Samples were subjected to 151 bp paired-end sequencing using a NovaSeqXPlus flow cell (Illumina, San Diego, CA) according to the manufacturer’s protocol. To determine quality of the data, reads were trimmed using Cutadapt v4.8 ^47^ and evaluated with FastQC v0.11.8 ^48^. Reads were mapped to the reference genome GRCm38 (ENSEMBL 109) using STAR v2.7.8a ^49^ and assigned count estimates to genes with RSEM v1.3.3 ^50^. Alignment options followed ENCODE standards for RNA-seq ^49^.

Unsupervised principal component analysis (PCA) based on variance-stabilized normalized reads and differential gene expression analysis was performed using R package DESeq2 v1.46.0 ^51^ with default parameters. To determine differentially expressed genes (DEGs), DESeq2 uses an empirical Bayes approach to integrate the dispersion and fold change estimates and tests the gene differential expression using the Wald test. The Wald test p values were adjusted for multiple testing using the procedure of Benjamini and Hochberg. Genes with an adjusted p value less than 0.05 and fold change greater than 1.5 or less than 0.67 were considered statistically significant. Differential expression data were visualized using volcano plots and the intersections of significant DEGs between each viral group and control were illustrated using upset plots. To compare the individual gene expression patterns of RV-A, RV-C and RSV with healthy controls, as well as the effects of all RV infections together with RSV and healthy controls, we performed sequencing analyses using two separate model designs to ensure normalization factors were accurately calculated in each comparison.

To categorize significantly upregulated and downregulated DEGs, enrichment of biological process gene ontology (GO) groups was performed using R package clusterProfiler v4.14.4 ^52^. Functional enrichment analysis of all DESeq2-normalized reads was preformed using GSEA v4.3.3 with dataset permutations and datasets available through MSigDB v2024.1.Hs ^53,54^. Normalized reads were ranked using the built-in Signal2Noise metric and a weighted enrichment statistic is reported.

### In silico analysis of cell type proportions

Cell-type proportions of the nasal swabs were estimated using the CIBERSORTx algorithm with B-mode batch correction ^55^. To maximize reference matrix sequencing depth, SmartSeq2-based single-cell data from the Human Lung Cell Atlas was used to generate a signature matrix for cell deconvolution ^56^. Cell types estimated by this reference cover all expected cell types in the upper airway by gentle nasal swabbing, excluding endothelial and stromal cell type to reduce error. To estimate relative eosinophil proportions between samples, we used the xCell algorithm with the standard xCell signatures (N=64) to analyze enrichment of eosinophilic marker genes ^57^.

### Data sharing

Processed count data and metadata will be uploaded to Gene Expression Omnibus upon publication. Due to donor privacy considerations, individual sequences will not be included.

## Results

### Sample inclusion and demographics

In general, children were excluded from study if they had a bacterial co-infection, viral co-infection (seven RSV samples) or asymptomatic viral infection (four control subjects). Eleven samples were excluded from RNA-seq analysis for poor RNA quality and ten samples were excluded due to unsuccessful viral typing. This left 46 samples from RV-infected patients, 12 samples from RSV-infected patients, and 6 samples from control subjects. Demographic data are shown in Table I. Thirty patients (51.7%) were infected with RV-C, 16 (27.6%) were infected with RV-A and 12 (20.7%) were infected with RSV. Patients with RSV were significantly younger, and control subjects significantly older, than patients with RV. There was no difference in gestational age at birth, sex, race, ethnicity, admission diagnosis, level of hospital care or day of sample collection between groups. A significantly higher proportion of patients with RV had an admission diagnosis of status asthmaticus, and a significantly higher proportion of patients with RSV had an admission diagnosis of bronchiolitis.

RNA transcripts in nasal swabs were measured by next generation sequencing and differences in gene expression calculated using DESeq2. Batch effects were found to be negligible, and the raw counts for technical replicates were combined using the collapseReplicates function for all subsequent analyses. To detect outliers, we performed unsupervised PCA based on variance-stabilized normalized reads using DESeq2. PCA verified that control samples clustered together (Figure 1A). In contrast, samples with viral infection showed substantial heterogeneity. No obvious outliers were identified.

**Figure 1.**
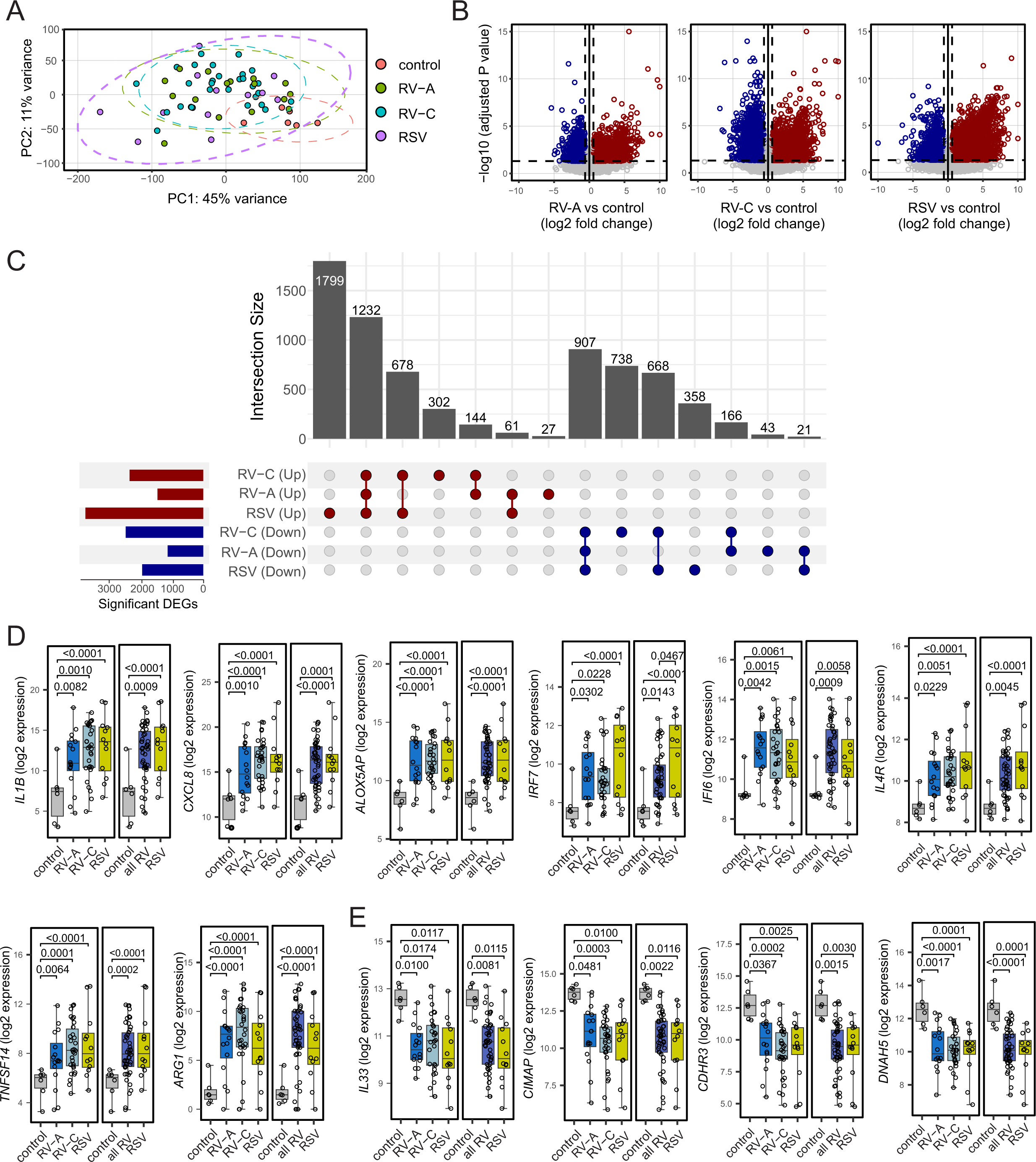
Effect of RV-A, RV-C and RSV on nasal epithelial gene expression. A. Unsupervised principal component analysis (PCA) based on variance-stabilized normalized reads was performed using DESeq2. Dotted ellipses depict a multivariate t-distribution around each group. B. Differential expression data were visualized using volcano plots. Color is based on the direction of regulation, with red signifying upregulation and blue signifying downregulation. C. The intersections of significant DEGs between each viral group and control were illustrated using upset plots. D. Selected DESeq2-normalized transcripts upregulated by RV-A, RV-C and RSV. E. Selected DESeq2-normalized transcripts downregulated by RV-A, RV-C and RSV. Shown are median, interquartile range (forming the box), and whiskers extending to the minimum and maximum values. Wald test p values were adjusted for multiple testing using the procedure of Benjamini and Hochberg. To compare the individual gene expression patterns of RV-A. RV-C and RSV with healthy controls, and to compare the effects of all RV infections together with RSV and healthy controls, we performed sequencing analyses using two separate model designs to ensure normalization factors were accurately calculated in each comparison.

### Genes regulated by all three viral species

RV-A induced differential expression of 2601 genes, 1464 of which were significantly upregulated and 1137 were significantly downregulated compared to controls (see Table E2 in the Online Repository). RV-C infection induced differential expression of 4,835 genes compared to controls, 2356 of which were upregulated and 2479 were downregulated (Table E2). Compared to controls, RSV induced differential expression of 5724 genes, 3770 of which were upregulated and 1954 were downregulated (Table E2). Volcano plots visualizing the direction, magnitude, and significance of changes in gene expression are shown in Figure 1B; an upset plot showing the numbers and intersections of differentially expressed genes between the three viruses are shown in Figure 1C. Twelve hundred and thirty-two genes were upregulated by each of the three viral species and included genes encoding interleukin (IL)-1β family cytokines, chemokines, tumor necrosis factor (TNF) family proteins, interferon response genes, and others. Selected genes upregulated by all three viruses are shown in Table III and Figure 1D. In particular, upregulated genes regulate granulocyte chemotaxis into the airways (*e.g., IL1A, IL1B, IL1RL2, IL36RN, CXCL2, CXCL8, CCL4L2, MCEMP1*), cysteinyl leukotriene metabolism (*ALOX5, ALOX5AP, LTB4R*), antiviral responses (*IRF7, IFITM2, IFI6, MEFV*), inflammasome activation (*NLRP3, IL18R*), while others are elevated in asthma (*IL4R, TNFSF14, ARG1, FCER1G*) or associated with asthma susceptibility (*IL1RL1, TNFAIP3, CCL3L1*).

**Table III.** Selected gene ontology biological process categories enriched by viral infection, as determined by GSEA.*

| *Gene Set* | *RV vs control* |  | *RSV vs control* |  | *RV vs RSV* |  |
| --- | --- | --- | --- | --- | --- | --- |
|  | *NES* | *FDR q* | *NES* | *FDR q* | *NES* | *FDR q* |
| regulation of innate immune response | 2.327 | **1.17E-04** | 2.753 | **0.00E+00** | -2.427 | **5.50E-04** |
| positive regulation of adaptive immune response | 1.924 | **7.69E-03** | 2.106 | **1.63E-03** | -2.299 | **5.48E-04** |
| pattern recognition receptor signaling pathway | 2.052 | **2.84E-03** | 2.540 | **6.15E-05** | -2.291 | **5.99E-04** |
| toll like receptor 2 signaling pathway | 2.087 | **2.23E-03** | 2.519 | **5.27E-05** | -2.200 | **7.33E-04** |
| toll like receptor 4 signaling pathway | 1.906 | **8.64E-03** | 1.906 | **8.64E-03** | -2.043 | **1.96E-03** |
| toll like receptor 3 signaling pathway | 1.420 | 1.05E-01 | 1.625 | **4.08E-02** | -1.709 | **2.36E-02** |
| regulation of natural killer cell activation | 1.592 | **4.92E-02** | 1.964 | **4.90E-03** | -2.218 | **5.47E-04** |
| granulocyte activation | 2.519 | **0.00E+00** | 2.131 | **1.26E-03** | -2.032 | **2.02E-03** |
| neutrophil chemotaxis | 2.504 | **0.00E+00** | 2.410 | **7.59E-05** | -2.016 | **2.21E-03** |
| myeloid dendritic cell activation | 2.017 | **3.60E-03** | 2.162 | **9.87E-04** | -1.947 | **3.87E-03** |
| macrophage activation | 2.317 | **1.12E-04** | 2.346 | **1.45E-04** | -1.693 | **2.58E-02** |
| mast cell activation | 2.102 | **1.97E-03** | 1.861 | **9.81E-03** | -1.517 | 6.87E-02 |
| alpha beta T cell activation | 2.044 | **3.08E-03** | 2.350 | **1.49E-04** | -2.261 | **8.01E-04** |
| T helper 17 type immune response | 1.763 | **2.08E-02** | 2.248 | **3.56E-04** | -2.264 | **7.04E-04** |
| positive regulation of Th1 type immune response | 1.711 | **2.71E-02** | 2.160 | **9.89E-04** | -1.900 | **5.87E-03** |
| type I interferon production | 1.744 | **2.29E-02** | 2.345 | **1.44E-04** | -1.989 | **2.73E-03** |
| positive regulation of type II interferon production | 1.857 | **1.17E-02** | 2.358 | **1.54E-04** | -1.997 | **2.54E-03** |
| inflammasome mediated signaling pathway | 1.761 | **2.09E-02** | 1.933 | **6.14E-03** | -1.734 | **2.02E-02** |
| interleukin 1 mediated signaling pathway | 1.955 | **5.92E-03** | 2.249 | **3.44E-04** | -2.194 | **7.49E-04** |
| interleukin 2 production | 1.915 | **8.22E-03** | 2.119 | **1.41E-03** | -1.815 | **1.15E-02** |
| positive regulation of interleukin 4 production | 1.813 | **1.59E-02** | 2.140 | **1.14E-03** | -1.665 | **3.01E-02** |
| response to interleukin 6 | 1.899 | **8.83E-03** | 2.202 | **6.93E-04** | -1.936 | **4.30E-03** |
| interleukin 8 production | 2.294 | **3.92E-04** | 2.486 | **4.10E-05** | -2.052 | **1.84E-03** |
| interleukin 10 production | 1.805 | **1.68E-02** | 2.416 | **7.74E-05** | -2.217 | **5.98E-04** |
| interleukin 12 production | 1.692 | **3.02E-02** | 2.279 | **2.61E-04** | -2.058 | **1.77E-03** |
| positive regulation of TNF superfamily production | 2.643 | **0.00E+00** | 2.888 | **0.00E+00** | -2.278 | **7.29E-04** |
| columnar cuboidal epithelial cell development | 2.108 | **1.84E-03** | 2.292 | **2.64E-04** | -1.341 | 1.59E-01 |
| epidermal cell differentiation | 1.750 | **2.20E-02** | 2.006 | **3.62E-03** | -1.914 | **5.28E-03** |
| positive regulation of EMT | 1.707 | **2.77E-02** | 1.657 | **3.47E-02** | -1.193 | 2.90E-01 |
| mucus secretion | 1.813 | **1.59E-02** | 1.455 | 9.20E-02 | 0.990 | 0.6494 |
| glycosylation | 1.969 | **3.48E-02** | -1.188 | 0.6888 | 1.969 | **3.48E-02** |

\*GSEA, gene set enrichment analysis; NES, normalized enrichment score; FDR, false discovery rate. Significant q values are denoted by **bold font**.

Nine hundred and seven genes were downregulated by RV-A, RV-C and RSV (Table E2). Selected genes are shown in Figure 1E. Downregulated transcripts encoded proteins associated with ciliated epithelial cells including the RV-C receptor *CDHR3, CIMAP,* and members of the cilia and flagella associated protein, dynein axoneme, Bardet-Biedl syndrome and intraflagellar transport families. *IL33* and *CCL14* were the only cytokine transcripts downregulated.

When we considered both RV species (RV-A and RV-C) together, RV infection significantly (adjusted p<0.05, fold change >1.5 or <0.67) regulated 4519 transcripts compared to controls (see Table E3 in the Online Repository). Volcano plots visualizing the direction, magnitude, and significance of changes in gene expression are shown in Figure 2A. A Venn diagram showing the numbers and intersections of differentially expressed genes between RV and RSV is shown in Figure 2B. Compared to uninfected controls, 1929 transcripts were upregulated by both RV and RSV.

**Figure 2.**
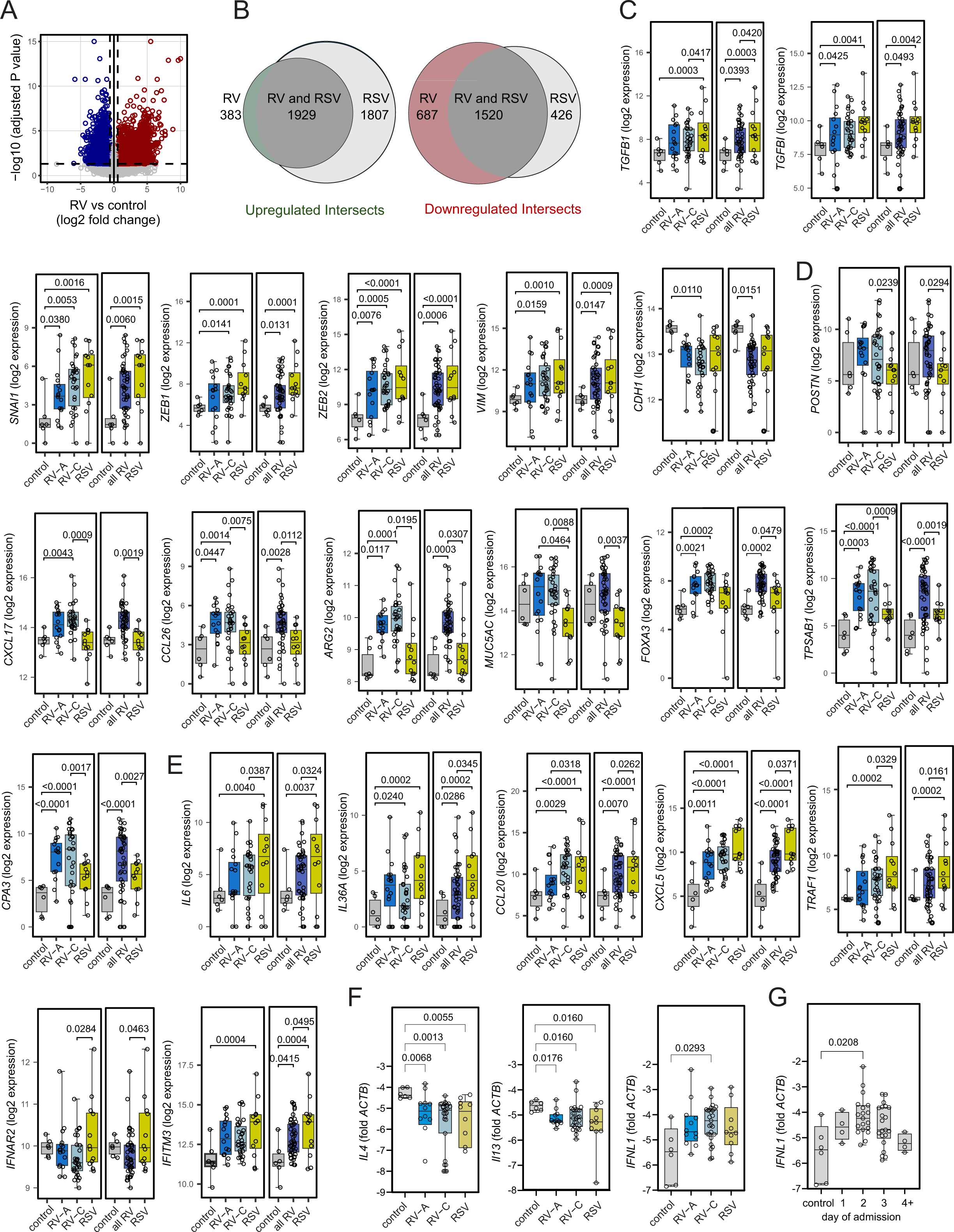
Combined effects of RV-A and RV-C on gene expression. A. Differential expression data were visualized using volcano plots. Color is based on the direction of regulation, with red signifying upregulation and blue signifying downregulation. B. Venn diagram showing the intersections of significant DEGs between RV and RSV. C. Box plots showing DESeq2-normalized EMT-related genes upregulated by RV-A, RV-C and RSV. D. Selected DESeq2-normalized transcripts upregulated by RV compared to RSV. E. Selected DESeq2-normalized transcripts upregulated by RSV compared to RV. Shown are median, interquartile range (forming the box), and whiskers extending to the minimum and maximum values. Wald test p values were adjusted for multiple testing using the procedure of Benjamini and Hochberg. F. qPCR data showing changes in IL-4, IL-13 and IFN-λ mRNA expression. Oligonucleotide primers used for qPCR are shown in Table 1. Statistical significance was assessed by the Kruskal-Wallis test for non-parametric, continuous variables.

We have shown in a cell culture model of injured/regenerating airway epithelium that RV infection induces EMT-like features ^42^. We therefore examined markers of EMT. Both RV and RSV significantly increased transcripts encoding transforming growth factor (TGF)-β1 and *TGFBI,* the EMT transcription factors *SNAI1* and *ZEB1* ^58^ and the extracellular matrix protein vimentin (Figure 2C). RV also decreased expression of the adherens junction protein E-cadherin.

### Differences in gene expression between viral species

Compared to samples from subjects with RSV, RV induced statistically significant upregulation of 207 genes (see Table E4 in the Online Repository). Upregulated genes included those involved in the regulation of eosinophilic inflammation (*CXCL17, CCL26, POSTN, ARG2),* mucus secretion (*MUC5AC, CLCA1, FOXA3, TFF1, TFF3*), and mast cell proteases *(CPA3, TPSAB1, TPSB2)* (Figure 2D). Compared to RV infections, RSV induced significantly greater upregulation of 674 genes including those involved in neutrophilic inflammation (*IL6, IL18, IL36A, TRAF1, CXCL5, CCL20*) and type 1 IFN response (*IRF7, IFNAR2, IFITM3*) (Figures 1D and 2E).

When we compared RV-C and RV-A directly, there were no statistically significant differences in gene expression. However, compared to controls there were 2386 genes that were significantly regulated by RV-C but not RV-A (980 upregulated, 1406 downregulated) (Table E2). Differential gene expression induced by RV-C infection was similar in magnitude to RSV (Figure 1C). Transcripts uniquely upregulated by RV-C included *CXCL17* (Figure 2D), *IFNL3, TPSD1* and *SPDEF*). Transcripts significantly regulated by RV-C and RSV but not RV-A included *ZEB1, VIM* (Figure 2C), *IL36A, CCL20* (Figure 2D), *TNF*, *IL36G, CCL23, IFIT2, IFIT3, MX2* and *OASL*. Transcripts downregulated by RV-C but not RV-A encoded proteins associated with ciliated epithelial cells (for example, *DNAH9*, *DNAI1, DNAI2* and *CFAP65*).

### Gene ontology analysis

To categorize significantly upregulated and downregulated DEGs, enrichment of biological process gene ontology (GO) groups was performed using R package clusterProfiler v4.14.4 ^52^. Among upregulated transcripts, 914 gene ontology (GO) groups were enriched after RV infection and 1527 GO groups enriched after RSV infection (see Figure E1 and Table E5 in the Online Repository). Relative to controls, GO groups representing activation, chemotaxis and/or differentiation of neutrophils, T cells, natural killer cells, dendritic cells and mast cells were significantly enriched for both RV and RSV. Both viruses showed enrichment of at least one GO group related to IL-1, IL-6, IL-8, IL-18, and TNF production.

Both RV and RSV showed enrichment of GOs related to epithelial cell proliferation, development and maturation. Both viruses showed enrichment of GOs related to T-helper 1 type immune response, T-helper 2 immune response and type 2 IFN response. RV and RSV each showed significant enrichment of GOs related to NF-κB, ERK, JAK/STAT signaling and Toll-like receptor (including MyD88-dependent) signaling. Only RV showed enrichment of genes associated with eosinophil migration and regulation of smooth muscle contraction. Only RSV showed enrichment of GOs related to inflammasome assembly, type 1 IFN response, and IL-2, IL-4 and IL-15 responses. Among downregulated transcripts, the two viruses showed nearly identical suppression of GOs related to cilium movement (not shown).

Functional enrichment analysis of all DESeq2-normalized reads was also performed using GSEA v4.3.3 with dataset permutations and molecular datasets available through MSigDB v2024.1.Hs ^53, 54^. This method allows a direct comparison of RV and RSV responses. In general, similar gene ontology groupings were enriched by both RV and RSV (Table III, Table E6 in the Online Repository). In nearly every case, enrichment was greater with RSV than RV. There were some new findings, however. In this analysis, RV infections enriched GO groups related to type 1 interferon production (albeit at a lower level than RSV). Also, enrichment of GOs related to mucus production was significantly greater for RV than RSV. In this analysis, neither virus significantly enriched GOs related to the T helper 2 type immune response.

### qPCR

Because transcripts encoding type 2 cytokines and IFNs were of low abundance, we performed qPCR for *IL4, IL5*, *IL13, IFNA1, IFNB1* and *IFNL1.* All three viral species significantly reduced expression of IL-4 mRNA, with an additional tendency towards reduced IL-13 (Figure 2F). qPCR showed that RV-C significantly induced IFN-λ mRNA expression.

While there was increased expression of IFN-stimulated genes (see Tables II, III), there was no detectable qPCR signal for *IFNA1* or *IFNB1* (not shown). We also examined the time of course of *IFNL1* mRNA expression. *IFNL1* appeared to peak 2 days after hospital admission (Figure 2F).

### Deconvolution of RNA-seq data

To examine the potential effects of viral infection on various airway cell populations, we performed computational deconvolution of RNA-seq data using the Human Lung Cell Atlas reference matrix ^56^. Focusing on nasal epithelial cells, RV-A, RV-C and RSV each significantly reduced the cell fraction of ciliated epithelial cells (Figure 3A). In addition, compared to RSV, RV-A and RV-C significantly increased the cell fraction of goblet cells. There was no effect on basal cell fraction. Turning towards inflammatory cells in the epithelium, all three viral species significantly increased the fraction of neutrophils and monocytes, while decreasing the fraction of naïve CD8+ T cells. There were trends towards an increase in CD4+ effector/memory cells and a reduction in dendritic cells. After RV-C infection, there was a significant reduction in macrophages and a trend towards increased mast cells/basophils (p=0.0732 vs control). Since the Human Lung Cell Atlas does not cover eosinophils, we also used the xCell 2.0 algorithm ^57^ for estimation of relative eosinophil populations (Figure 3B). There was substantial heterogeneity in the eosinophil response to viral infection and there were no differences between the three viral species.

**Figure 3.**
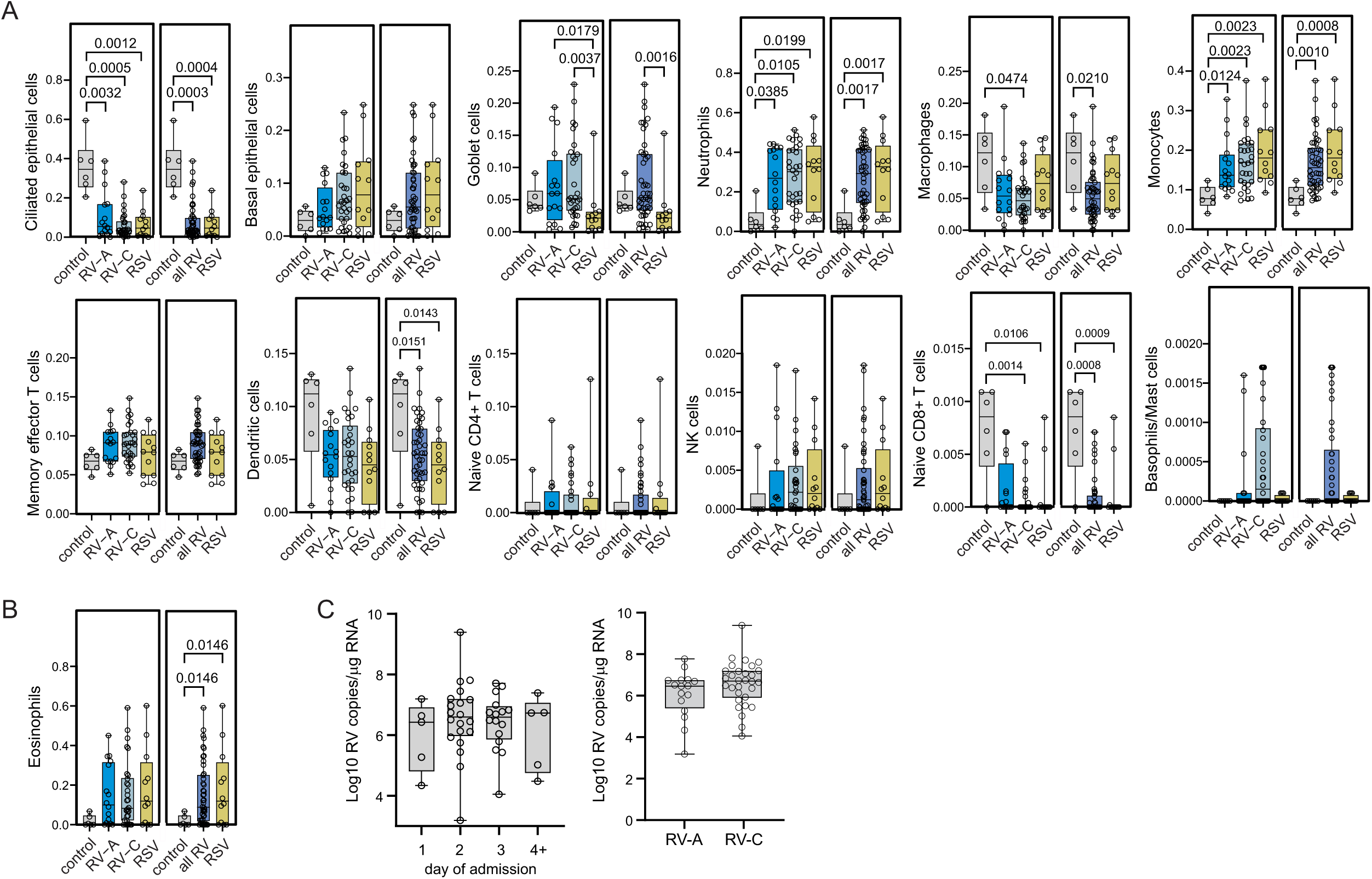
Computation deconvolution of differentially expressed genes showing changes in cell proportion. A. Cell-type proportions of the nasal swabs were estimated using the CIBERSORTx algorithm and signature matrices from the Human Lung Cell Atlas. B. To estimate relative eosinophil proportions between samples, we used the xCell algorithm. C. Viral copy number for RV infections was measured by qPCR using pan-RV primers. Data were normalized to 1 µg total RNA. Shown are median, interquartile range (forming the box), and whiskers extending to the minimum and maximum values. Statistical significance was assessed by the Kruskal-Wallis test for non-parametric, continuous variables.

### RV copy number

We measured viral copy number for RV infections by qPCR using pan-RV primers. Data were normalized to 1 µg total RNA (Figure 3C). RV copy number tended to peak on day 2 after admission. There was no difference in viral copy number between RV-A and RV-C infections. However, there were many significant associations between viral copy number and gene expression. After RV-A infection, 376 genes had a strongly positive correlation with viral copy number (Pearson correlation coefficient ≥ 0.5), and 684 additional genes had a moderate positive correlation (Pearson correlation coefficient between ± 0.40 and ± 0.49). After RV-C infection, 240 genes had a strongly positive correlation with viral copy number (Pearson correlation coefficient ≥ 0.5), and 693 additional genes had a moderate positive correlation (Pearson correlation coefficient between ± 0.40 and ± 0.49). Genes that were upregulated with viral infection and had a positive correlation with RV-C copy number included the chemokine *CXCL13*; interferons *IFNL1, IFNL2* and *IFNL3;* the viral response genes *RIGI, IFIH1, IFI3, IFI16, IFI44, IFI44L, IFIT1, IFIT2, IFIT3, IFIT5, AIM2, MX1, OAS2, OAS3 and OASL;* and the mast cell proteases *CPA3, TPSAB1* and *TPSB2* (Figure 4A). Transcripts that were downregulated with viral infection but has a positive correlation with RV-C copy number included *IL33, IFNG,* the chemokines *CXCL9, CXCL10, CCL5, CXCL14, CCL15 and CCL5*; and the RV-C receptor *CDHR3* (moderately positive) (Figure 4B).

**Figure 4.**
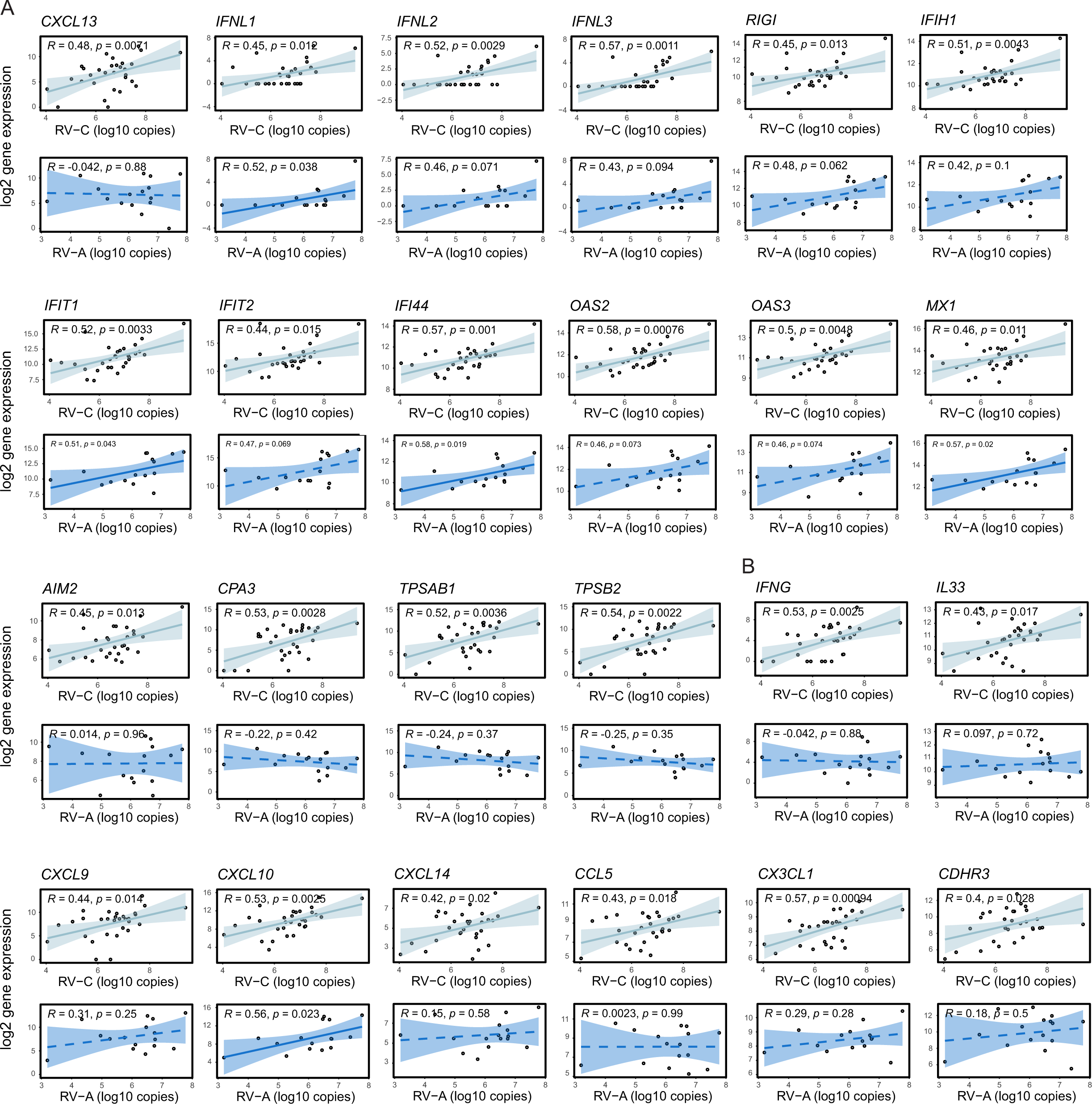
Correlation of viral copy number and gene expression. Viral copy number was measured by qPCR and expressed as log10 copies relative to total RNA. Gene expression was expressed as log2 normalized transcripts. A. Transcripts that were upregulated with RV infection compared to controls. B. Transcripts that were downregulated with RV infection. The significance of correlations between viral copy and gene expression was determined using the Pearson correlation coefficient. Solid trendlines depict significant correlation, while dashed lines depict insignificant correlation.

## Discussion

We measured respiratory tract epithelial gene expression in hospitalized children with RV and RSV infections. We found that evidence of a robust type 1 inflammatory response, with enrichment of transcripts encoding activation, chemotaxis and/or differentiation of neutrophils, eosinophils, T cells, natural killer cells, γδ T cells and dendritic cells. Especially predominant was upregulation of genes regulating granulocyte chemotaxis. Samples from RV and RSV infections showed enrichment of at least one GO group related to IL-1, IL-6, IL-8, IL-18, and TNF production. In addition to genes encoding CXC chemokines and TNF family proteins, we found significant overexpression of transcripts encoding IL-1 family cytokines including IL-1α, IL-1β and, for RV-C and RSV, IL-36α and IL-36γ. IL-36 cytokines are inactive as full-length proteins and require activation by soluble neutrophil proteases ^59^. *IL36G* has recently been shown to drive neutrophilic inflammation in COPD ^60^ but has not yet been shown to be induced by viral infection in humans. Consistent with the overexpression of neutrophil chemoattractants, computational deconvolution showed an increased in nasal neutrophils after viral infection.

While RV and RSV elicited a robust type 1 immune response, the effect of respiratory viral infection on type 2 inflammation was mixed. Analysis of DEGs showed enrichment of the “regulation of T-helper 2 cell differentiation” gene ontology grouping by both RV and RSV. On the other hand, GSEA did not show enrichment of transcripts related to the Th2 response, and we found reductions in mRNA expression of IL-33 (by DESeq2), IL-4 and IL-13 (by qPCR).

Following viral infection, prioritization of antiviral immunity may lead to suppression of the type 2 immune response. RSV ^61^ and IFNs ^62^ have each been shown to repress leukocyte IL-4 expression, and children with IRF7^hi^ nasal responses to respiratory viral infection showed reduced *IL33* ^39^.

RV and RSV also increased expression of *ALOX5AP, ALOX5* and *LTB4R*, implicating cysteinyl leukotrienes (CysLTs) in the response to viral infection. CysLTs have been shown to be increased in the nasopharygneal secretions of children with virus-induced wheezing ^63^. A genetic variant in *ALOX5* has been associated with reduced lung function in children with poorly controlled asthma ^64^. CysLTs are produced by mast cells, eosinophils and basophils, each of which were identified in our nasal swab samples by computational deconvolution. In the mouse, CysLTs are also produced by airway tuft cells, a rare epithelial cell responsible for allergen-^65^ and viral-induced eosinophilic inflammation ^66^. Since human tuft cells express *ALOX5AP* ^67^, it is therefore conceivable that tuft cells are a source of *ALOX5AP* and *ALOX5* in our samples.

RV and RSV infections each upregulated genes associated with epithelial remodeling and EMT including *TGFB, TGFB1, SNAI1, ZEB1, ZEB2* and *VIM.* Airway epithelial repair begins with EMT, a process in which non-motile epithelial cells gain motility, migratory, and invasive properties. Once the epithelial barrier is re-established, epithelial cells within the basal compartment undergo ciliogenesis or differentiate into secretory cells to re-establish a pseudostratified mucociliary epithelium. TGF-β-induced EMT is increased in airway epithelial cells from patients with asthma ^68^. Finally, we have shown in a cell culture model of injured/regenerating airway epithelium that RV infection induces EMT-like features such as reduced transepithelial resistance and reduced expression of occludin and E-cadherin ^42^.

Together these data suggest the possibility that the airway epithelium of children with respiratory viral infections undergo epithelial remodeling or partial EMT.

RV and RSV infections each downregulated expression of genes responsible for ciliary organization and assembly. Previous *in vitro* studies have revealed a common transcriptomic signature shared by cytotoxic respiratory viruses, including RSV, which includes the global downregulation of cilium-related genes ^69, 70^. In a study of RV-infected cells, RV-C but not RV-B downregulated expression of cilia-related genes ^71^. In addition to the suppression of gene expression, the observed reduction in transcripts regulating ciliary organization and assembly could also be due to a reduction in the number of ciliated epithelial cells. Reduction of ciliated epithelial cells has been previously shown for cell cultures infected with RSV ^70^. In our study, we confirmed using computational deconvolution of RNA-seq data that RV-A, RV-C and RSV each significantly reduce the cell fraction of ciliated epithelial cells *in vivo*.

Focusing on differences in the nasal epithelial transcriptome between viral species, compared to RSV, RV induced statistically significant upregulation of genes involved in the regulation of eosinophilic inflammation (*CXCL17, CCL26, POSTN, ARG2),* mucus secretion (*MUC5AC, CLCA1, FOXA3*), and mast cell proteases *(CPA3, TPSAB1, TPSB2).* Accordingly, RV infection was associated with an increase in nasal goblet cells. In addition, RV-C (but not RV-A) induced an increase in the fraction of nasal mast cells. The expression of mast cell proteases and chemokines responsible for mast cell migration (*CX3CL1, CXCL10*) ^72–74^ was correlated with RV-C copy number, evidence of mast cell involvement in RV-C-induced respiratory illnesses. While mast cells were once confined by many workers to a secondary role in asthma exacerbations, they are now recognized as both first-line sentinels of the innate immune system and important modulators of downstream allergic responses. In patients with severe asthma, basophils and mast cells were found to be the most predominant producers of IL-4 and IL-13 in the airways ^75^. Expression of *CPA3, TPSAB1* and *TPSB2* is increased in the airway epithelium of asthmatic patients ^44^. Mast cells and mast cell proteases are found within the airway smooth muscle, epithelium and subepithelium of both high Th2 and low Th2 asthma, and their numbers correlate with sputum Th2 cytokines ^43, 76–81^. The presence of mast cells in the airway smooth muscle suggests that these cells are critical determinants of airways hyperresponsiveness. Besides the IgE receptor FCεR1, mast cells express Toll-like receptors 1-9 and express type 1 IFNs upon TLR3 stimulation ^82^. Finally, nasal brush samples taken from adult controls and allergic subjects during and after upper respiratory tract viral infections (including RV) showed increased tryptase-positive mast cells in allergic subjects ^83^. Taken together, these data suggest that mast cells may play an important functional role in response to respiratory viral infection in hospitalized children.

In contrast to our previous community-based study ^84^, we observed more infections with RV-C than RV-A. RV-C tended to be a stronger regulator of epithelial gene expression than RV-A. Of the three viral species, only RV-C infection significantly induced IFN-λ1 mRNA expression. In asthmatic children with RV infections, increased IFN-λ1 protein levels are associated with wheezing and exacerbation severity ^85^. Aside from the increase in IFN-λ by RV-C, we did not find a significant increases in *IFN* transcripts after viral infection. The lack of viral transcripts might have been due to low enrichment of immune cells in the airway epithelium.

However, we found that RV infection significantly increased the expression of a number of IFN response genes, many of which correlated with viral copy number. Also, our GSEA showed enrichment of transcripts encoding a type 1 IFN response. At first blush, these data would seem to suggest that children hospitalized with viral infections, some of whom have asthma, had an appropriate IFN response to viral infection, though we do not have a control group of infected healthy children with whom to compare. Finally, it should be noted that, compared to controls, RSV proved to be a stronger regulator of differential epithelial cell gene expression than RV, including genes stimulated by type 1 IFNs.

We would like to point out the limitations to our study. As noted above, we did not have a control group of healthy children with respiratory viral infections with whom to compare results. We were unable to recruit many control subjects, and controls were older than hospitalized patients with viral infection. We did not obtain bronchial epithelial samples for analysis, but previous studies have demonstrated concordance in gene expression between upper and lower airway samples ^35^.

In conclusion, we measured respiratory tract epithelial gene expression in hospitalized children with RV and RSV infections, employing computational deconvolution to assess cell fractions and correlating gene expression with viral copy number. RV and RSV each induced a robust type 1 inflammatory response with enrichment of transcripts encoding genes regulating leukocyte chemotaxis. However, there were some differences between viral species. RSV proved to be a stronger stimulus of differential epithelial cell gene expression than RV, and RV-C was a stronger regulator of gene expression than RV-A. RV induced significant upregulation of genes regulating eosinophilic inflammation, mucus secretion and mast cell proteases. RV-C infections, but not those caused by RV-A or RSV, were associated with an increase in mast cells in the nasal epithelium, as well as increased IFN-λ. Together, these data provide insight into the mechanisms by which RV, and in particular, RV-C, trigger respiratory exacerbations.

## Abbreviations

CDHR3: cadherin-related family member 3
CystLT: cysteinyl leukotriene
DEG: differentially expressed gene
EMT: epithelial-to-mesenchymal transition
GO: gene ontology
ICAM-1: intercellular adhesion molecule-1
IFN: interferon
IL-1: interleukin-1
LDL-R: low-density lipoprotein receptor
PCA: principal component analysis
PCR: polymerase chain reaction
RIN: RNA integrity number
RNA-Seq: RNA sequencing
RV: rhinovirus
TGF-β1: transforming growth factor-beta
TLR: Toll-like transcription factor
TNF: tumor necrosis factor

---

**Supplemental Figure E1.**
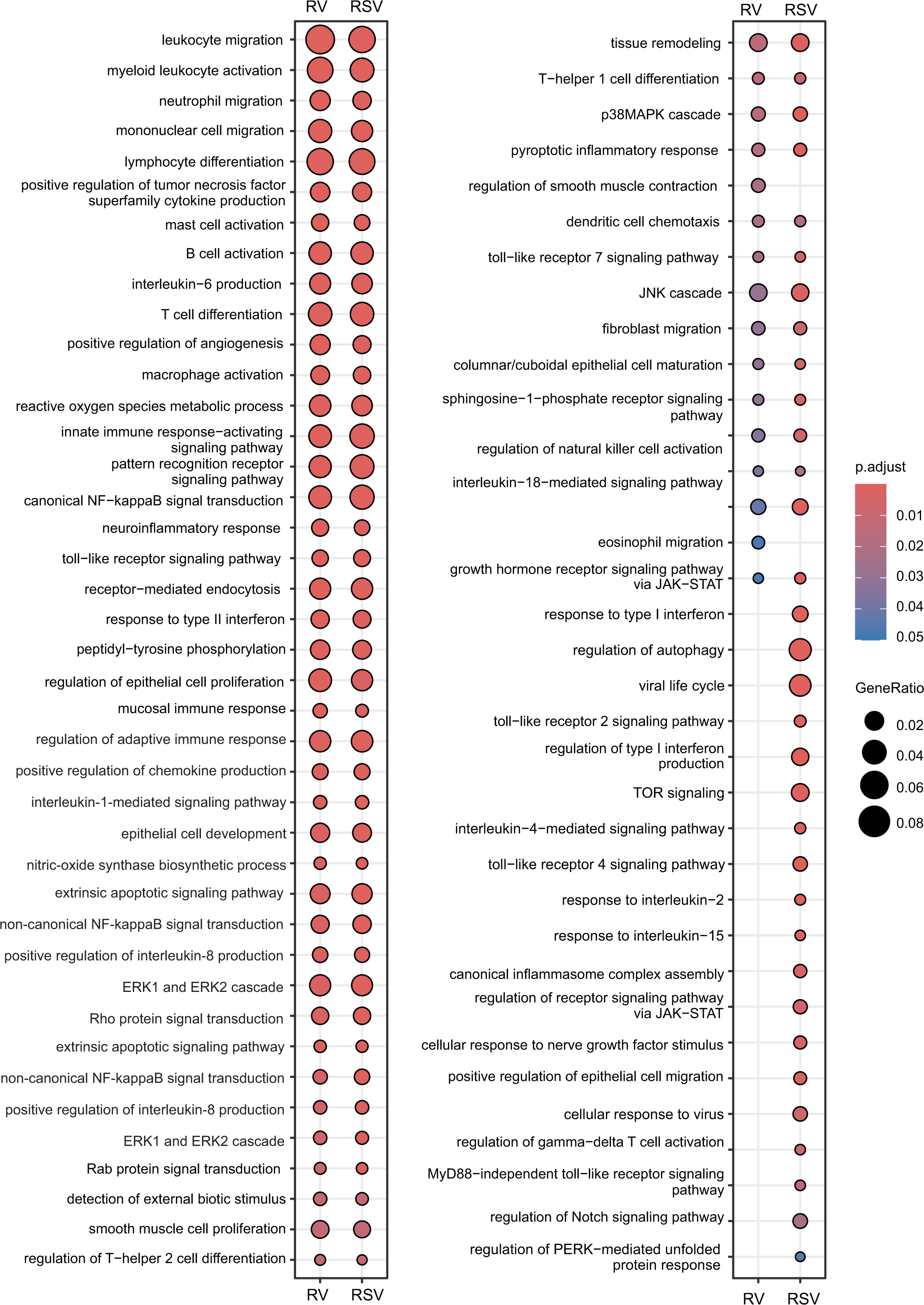
Enrichment of biological process gene ontology groups by RV and RSV. To categorize significantly upregulated and downregulated DEGs, enrichment of biological process gene ontology (GO) groups was performed using R package clusterProfiler v4.14.4. P values were adjusted for multiple testing using the procedure of Benjamini and Hochberg. The color of the circle indicates the adjusted p value for the degree of enrichment. In this scale, red represents a smaller p value and a higher enrichment degree. The size of circle represents gene ratio.

