## Supplemental Table 1 for "Upper Airway Gene Expression in Hospitalized Children with Rhinovirus-induced Respiratory Illnesses"

**Table E1.** PCR primers used.

*Transcript	Primers	Sequence*

RV	forward	5’-GTC CTC CGG CCC CTG AAT G-3’

reverse	5’-GAA ACA CGG ACA CCC AAA GTA G-3’

β-actin	forward	5’-CAC CAT TGG CAA TGA GCG GTT C-3’

reverse	5’-AGG TCT TTG CGG ATG TCC ACG T-3’

IFN-α1	forward	5’-ACT CAT ACA CCA GGT CAC GC-3’

reverse	5’-TGG TCA TAG TTA TAG CAG GGG TG-3’

IFN-β1	forward	5’-AAC ATG ACC AAC AAG TGT CTC C-3’

reverse	5’-GGA ATC CAA GCA AGT TGT AGC-3’

IFN-λ1	forward	5’-GGA CGC CTT GGA AGA GTC ACT-3’

reverse	5’-AGA AGC CTC AGG TCC CAA TTC-3’

IL-4	forward	5’-CCA ACT GCT TCC CCC TCT G-3’

reverse	5’-TCT GTT ACG GTC AAC TCG GTG-3’

IL-5	forward	5’-GGA ATA GGC ACA CTG GAG AGT C-3’

reverse	5’-CTC TCC GTC TTT CTT CTC CAC AC-3’

IL-13	forward	5’-CAC GGT CAT TGC TCT CAC TT-3’

reverse	5’-TGG TTC TGG GTG ATG TTG AC-3’
